# Engineered Extracellular Vesicles Enriched with the miR-214/199a Cluster Enhance the Efficacy of Chemotherapy for Ovarian Cancer

**DOI:** 10.1101/2025.07.18.665422

**Authors:** Weida Wang, Ayesha Alvero, Yi Qin, Mingjin Wang, Alexandra Fox, Yanfeng Li, Michael Millman, Amy Kemper, Gil Mor, Xian Shuang Liu, Michael Chopp, Zheng Gang Zhang, Yi Zhang

## Abstract

Recurrent ovarian cancer (OC) remains a major cause of mortality due to chemoresistance and metastasis. Epigenetic dysfunction, particularly through altered microRNA (miRNA) expression, contributes to disease progression. Targeting these molecular aberrations is critical to prevent recurrence, limit metastasis and improve patient outcomes. Here, we identify the miR-214-3p/miR-199a-5p cluster as a stage-associated, tumor-suppressive network that is lost in recurrent and chemoresistant OC, but can be restored using engineered small extracellular vesicles enriched with this cluster (m214-sEVs). Using a clinically relevant mouse model that mimics spontaneous OC relapse following first-line platinum-based chemotherapy, we showed that m214-sEVs were internalized by OC cells and the OC niche fibroblasts via clathrin-mediated endocytosis, resulting in the elevation of miR-214-3p/miR-199a-5p and the downregulation of chemoresistance-associated genes, including toll-like receptor 4 (TLR4), β-catenin, and the soluble N-ethylmaleimide-sensitive factor attachment protein receptor (SNARE) protein YKT6. Moreover, secondary tumor-derived sEVs (t-sEVs) released by OC and niche cells that internalized m214-sEVs reduced pro-metastatic proteins, such as integrin β1 and matrix metalloproteinase 9 (MMP9), in their cargo and limited their capacity to promote invasion and resistance. In vitro, YKT6 overexpression in ovarian cancer stem cells (OCSCs) attenuated the effect of m214-sEVs on sensitizing carboplatin to block OCSC migration. These findings demonstrate that engineered m214-sEVs designed to restore clinically lost tumor-suppressive miRNAs can concurrently reverse chemoresistance and reprogram tumor-derived EV communication by targeting oncogenic networks.

**Statement of Significance:** Engineered small extracellular vesicles delivering miR-214-3p/miR-199a-5p overcome chemoresistance and inhibit recurrence in ovarian cancer by targeting oncogenic networks and reprogramming tumor-derived extracellular vesicle communication within the tumor microenvironment.

## Introduction

Ovarian cancer (OC) is a highly heterogeneous disease driven by diverse cellular phenotypes that contribute to its complexity and high recurrence rate [1, 2]. Recurrent OC remains largely incurable due to resistance to standard therapies and aggressive metastatic behavior, with second-line treatments rarely yielding durable responses[3, 4]. Compared to primary tumors, recurrent OC is enriched in ovarian cancer stem cells (OCSCs) and cancer-associated fibroblasts, cell populations that promote therapeutic resistance and poor clinical outcomes [5, 6].

MicroRNA (miRNA) dysregulation is a hallmark of epigenetic dysfunction in cancer, orchestrating gene networks that govern OC progression and treatment response [7, 8]. Aberrant miRNA expression contributes to tumor heterogeneity and exhibits cell-type-specific and stage-dependent patterns aligned with disease advancement [9, 10]. While oncogenic miRNAs enhance tumor aggressiveness, loss of tumor-suppressive miRNAs is frequently associated with advanced disease and poor prognosis [11], Although some studies report miR-214 upregulation [12, 13], approximately 45% of patients with advanced OC and 60% of those with high-grade serous ovarian carcinoma (HGSOC) display marked downregulation of miR-214 and its cluster partner miR-199a[14]. Functionally, miR-214 overexpression suppresses OC growth by targeting phosphatase and tensin homolog deleted on chromosome 10 (PTEN) and semaphorin signaling[12], and our previous work showed that loss of the miR-214-3p/199a-5p cluster in OCSCs promotes tumor progression and chemoresistance[15, 16]. Together, these findings suggest the miR-214/199a cluster as a clinically significant tumor-suppressive network whose loss drives recurrence and drug resistance.

Small extracellular vesicles (sEVs) are membrane-bound nanovesicles that mediate intercellular communication by transferring bioactive cargo, including miRNAs [17, 18]. Compared with synthetic delivery systems, sEVs provide efficient, biocompatible, and cell-specific delivery [19, 20]. Within the tumor microenvironment (TME), sEVs regulate key processes such as chemoresistance, metastasis, and immune evasion[7, 21]. While tumor-derived sEVs often promote malignancy[22, 23], sEVs released from non-malignant cells, such as mesenchymal stromal cells (MSCs)[24], normal fibroblasts[25], and cerebral endothelial cells (CECs)[9, 26], can exhibit tumor-suppressive effects. Emerging evidence further suggests that therapeutic sEVs can reprogram the release and composition of secondary EVs from tumor and stromal cells, amplifying their therapeutic impact[27].

Our previous work demonstrated that CEC-derived sEVs (CEC-sEVs) mitigate chemotherapy-induced peripheral neuropathy and enhance the cytotoxicity of OC treatments by downregulating genes linked to neurotoxicity and tumor progression[9]. Building on these findings, the present study examined the clinical expression pattern of the miR-214-3p/miR-199a-5p cluster during OC progression and tested the hypothesis that engineered CEC-sEVs enriched with the miR-214-3p/miR-199a-5p cluster (m214-sEVs) can enhance chemotherapy efficacy and prevent recurrence in a clinically relevant mouse model of recurrent OC.

## Results

### Loss of the miR-214-3p/miR-199a-5p Cluster Correlates with Chemoresistance and Recurrence in OC

We previously identified miR-214-3p and its cluster partner miR-199a-5p as among the most downregulated miRNAs in chemoresistant OCSCs [15]. Analysis of The Cancer Genome Atlas (TCGA) dataset confirmed significant reductions in both miRNAs in HGSOC patients with grade 3/4 tumors compared to those with grade 1/2 disease, with a more pronounced decrease in miR-199a-5p (**Fig. 1A**). Prognostic survival analysis of the tumor-free HGSOC patient subset, defined as patients without residual disease or recurrence at last follow-up, showed that higher expression levels of these miRNAs were strongly associated with prolonged tumor-free survival (**Fig. 1B**). Furthermore, in situ hybridization (ISH) of recurrent HGSOC specimens demonstrated minimal or undetectable expression of both miRNAs in tumor epithelial cells, but robust expression in adjacent stromal compartments (**Supplemental Fig. 1**), indicating a cell-type-specific loss of this tumor suppressive miRNA cluster during OC progression.

**Figure 1.**
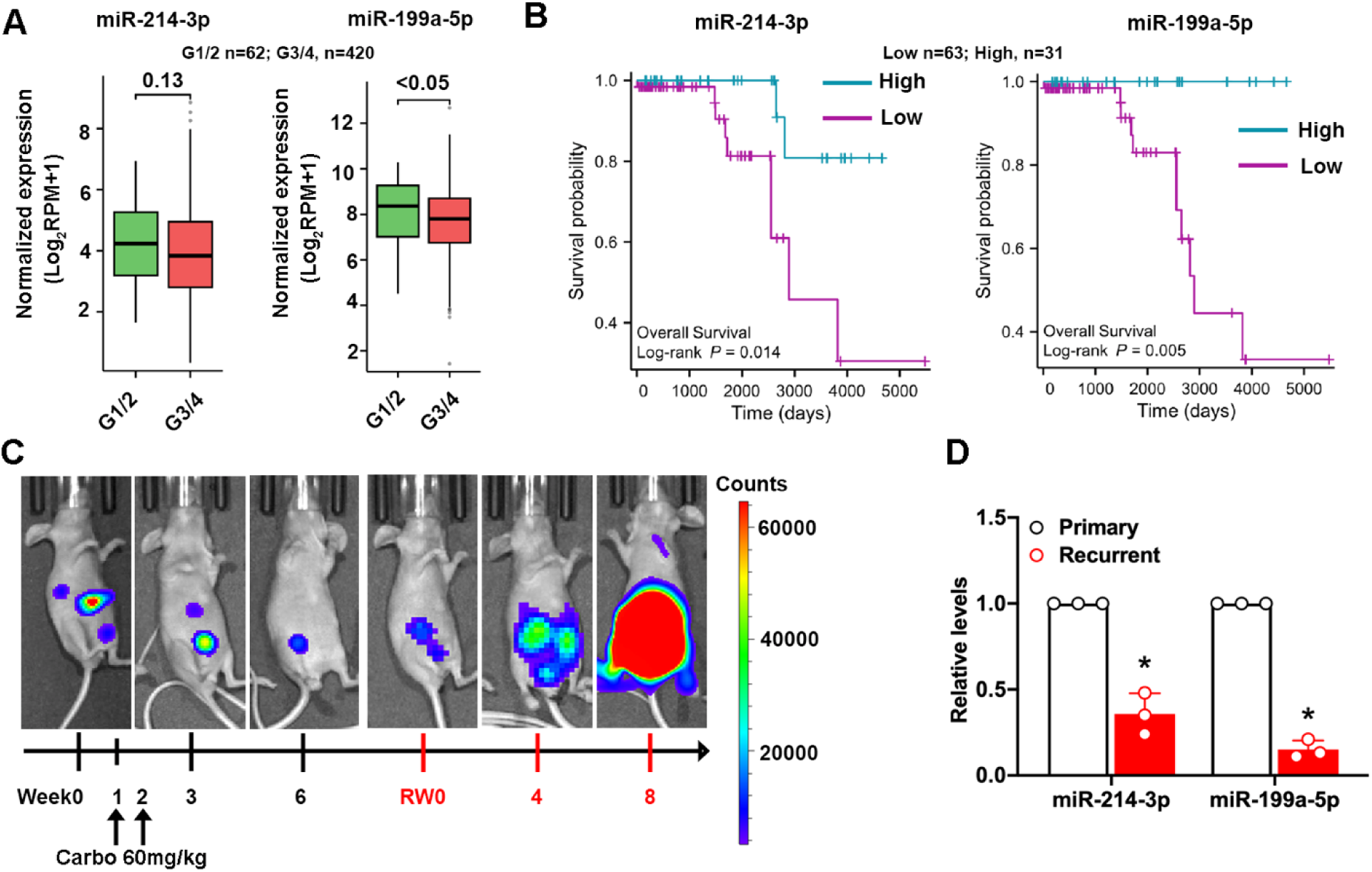
Tumor Recurrence in a Xenograft Model of OC and Downregulation of miR-214-3p and miR-199a-5p in recurrent tumors. **(A)** Expression levels of miR-214-3p and miR-199a-5p in human HGSOC samples from the TCGA-OV dataset analyzed by histological grade (G1+G2, n = 62; G3+G4, n = 420). (**B**) Kaplan–Meier survival analysis of overall survival based on miRNA expression levels. Tumor-free patients were dichotomized into high-(n=31) and low-expression groups (n=63) using the optimal cut-off. (**C**) Representative BLI of mice bearing intraperitoneal OVCAR3/luc xenografts treated with carboplatin (60 mg/kg) at weeks 1 (W1) and 2 (W2) post-implantation. Tumor recurrence was first detected and designated as recurrence week 0 (RW0) and progressed through recurrence weeks 4 (RW4) and 8 (RW8). The color scale reflects photon flux intensity (photons/sec/cm²). **(D)** Quantitative RT-PCR analysis of miR-214-3p and miR-199a-5p expression in primary tumors (W2) compared to recurrent tumors (RW4). Data are presented as mean ± SEM. *p < 0.05 versus primary group, determined by one-way ANOVA.

To determine whether this pattern is recapitulated in vivo, mice bearing OVCAR3/luc xenografts [9] with carboplatin at a clinically relevant regimen (60 mg/kg, tail-vein on days 7 and 14 post-implantation) [28, 29] (**Fig.1C**). Six weeks after treatment, tumors were undetectable by bioluminescence imaging (BLI; total flux < 5×10^5^ p/s/cm^2^). However, approximately 40% of mice developed recurrent tumors (flux > 1×10^6^ p/s/cm^2^) within eight weeks post-treatment (recurrent week 0, RW0; **Fig. 1C**), leading to 100% mortality by RW8. Recurrent lesions exhibited markedly reduced miR-214-3p and miR-199a-5p expression compared with primary xenografts (**Fig. 1D**). Together, these findings demonstrate that miR-214-3p and miR-199a-5p are downregulated in advanced and recurrent OC across patient and animal models, supporting their roles as tumor-suppressive miRNAs in disease progression.

### Engineered miR-214-3p/miR-199a-5p-enriched sEVs Enhance the Therapeutic Efficacy of Carboplatin in Recurrent OC

Building on our previous findings that CEC-derived sEVs enhance chemotherapy efficacy in animal models [9], we investigated whether engineered sEVs enriched with miR-214-3p/miR-199a-5p (m214-sEVs) could similarly improve chemotherapy response in recurrent OC. We generated m214-sEVs by transducing human CECs with a lentiviral vector encoding miR-214-3p as previous described [30]. The transduced CECs maintained typical endothelial morphology and ZO-1 expression, indicating no detectable alteration in cell phenotype (**Fig. 2A**). sEVs were isolated from conditioned media and characterized following Minimal Information for Studies of Extracellular Vesicles (MISEV) 2018/2023 guidelines[31, 32]. Transmission electron microscopy (TEM), cryo-electron microscopy, and nanoparticle tracking analysis (NTA) confirmed comparable sEV morphology and size (**Fig. 2B-D**), while Western blot and ExoView verified comparable tetraspanin enrichment (CD63, CD81, CD9) and absence of the endoplasmic reticulum (ER) marker calnexin cross m214-sEVs, sEVs from scramble-transfected CECs (scra-sEVs), and naïve CEC-sEVs (**Fig. 2E-F, Supplemental Fig. 2**). These findings indicate that lentiviral transduction did not alter vesicle morphology or yield [30]. However, qRT-PCR analysis showed an ∼11-fold increase of miR-214-3p and ∼6-fold increase of miR-199a-5p in m214-sEVs compared with scra-sEVs and CEC-sEVs, with no change in unrelated miRNAs (miR-15b-5p, miR-16-2) (**Fig. 2G**). Proteomic analysis showed that the overall protein composition of m214-sEVs was comparable to that of naïve CEC-sEVs (**Supplemental Excel 1**). The most enriched Gene Ontology (GO) terms included cadherin binding and RNA binding processes, which are functions associated with sEV-recipient cell interactions and cargo sorting, respectively (**Supplemental Fig. 3, Supplemental Excel 2**). These results confirm that m214-sEVs selectively enrich therapeutic miRNAs without altering the core protein cargo.

**Figure 2:**
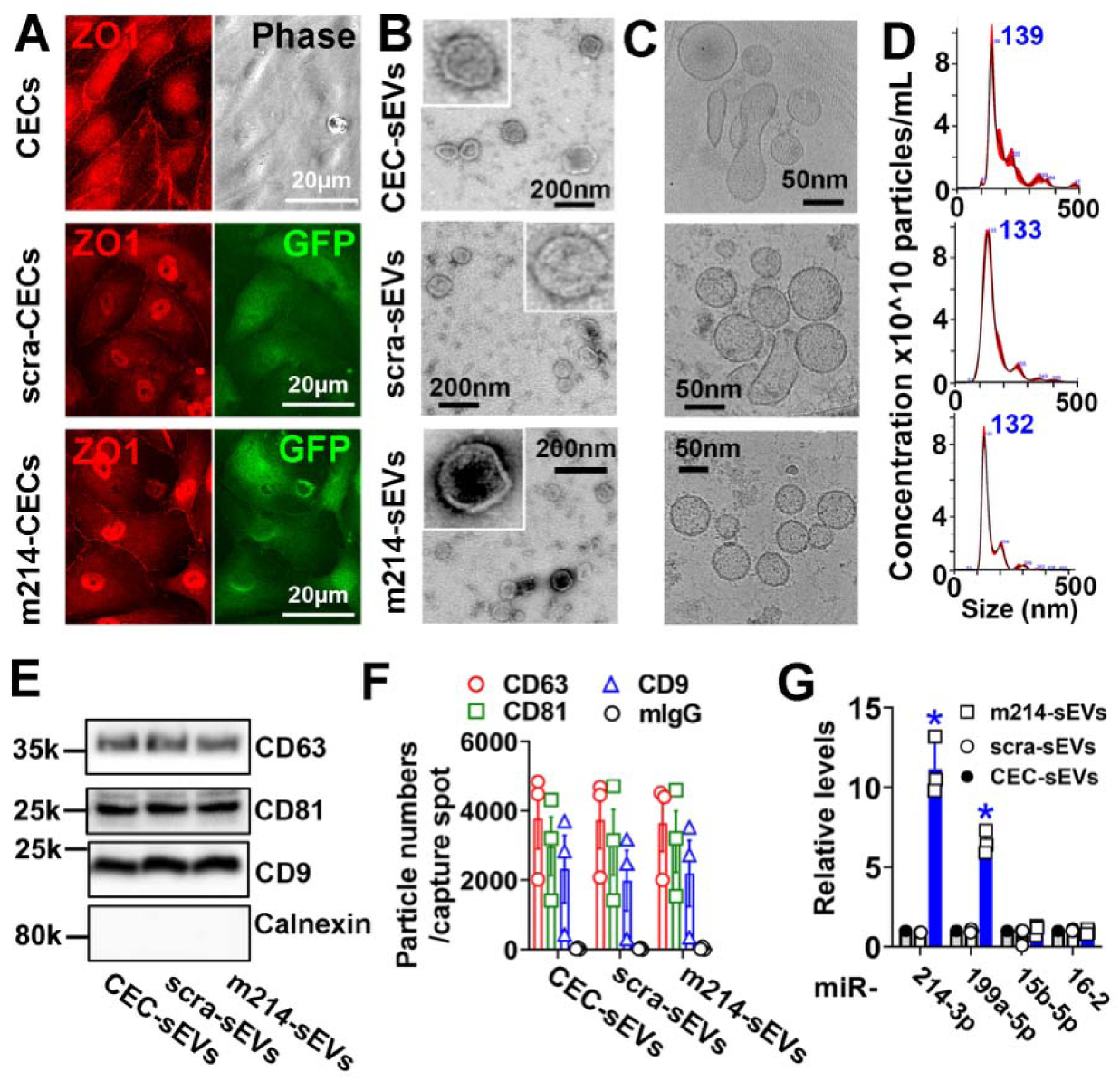
Characterization of Engineered CEC-Derived sEVs Enriched with miR-214-3p. (**A**) Representative fluorescence images of naïve CECs (with phase image), and CECs transduced with either miR-214-3p (m214-CECs) or scrambled control (scra-CECs), showing robust GFP expression and preserved endothelial morphology, as indicated by the tight junction marker ZO1. (**B, C**) Representative TEM with enlarged views and cryo-EM images show isolated sEVs from CECs (CEC-sEVs), m214-CECs (m214-sEVs) and scra-CECs (scra-sEVs). (**D**) NTA results show comparable size distributions of CEC-sEVs, m214-sEVs and scra-sEVs. (**E**) Western blot results demonstrate the presence of tetraspanin markers (CD63, CD81, CD9) and the absence of the ER marker calnexin, indicating the purity of vesicle preparations. (**F**) ExoView-based quantification of CD63⁺, CD81⁺, and CD9⁺ particle counts across EV preparations. (**G**) Quantitative RT-PCR analysis of miR-214-3p, miR-199a-5p, miR-15b-3p, and miR-16-2 in the different EV samples. *p < 0.05 vs. scra-sEVs, assessed by one-way ANOVA.

We next evaluated the therapeutic efficacy of m214-sEVs *in vivo*. As our prior studies showed that CEC-sEVs alone do not alter OC progression[9], we assessed the efficacy of m214-sEVs in a recurrent OC model in combination with chemotherapy. Mice bearing recurrent OVCAR3/luc tumors were randomized to receive PBS, carboplatin alone, carboplatin plus scra-sEVs, or carboplatin plus m214-sEVs (**Fig. 3A**). Carboplatin (60 mg/kg) was given on RW1 and RW2, and sEVs (1×10^11^ particles, intraperitoneally, i.p.) every other day for six weeks. The PBS-treated group exhibited rapid tumor progression, with all animals reaching experimental endpoint by RW6 (**Fig. 3B-D**). Treatment with carboplatin alone or in combination with scra-sEVs significantly decreased tumor growth compared to PBS control, but still, all animals in these groups reached the endpoint by RW10 (**Fig. 3B-D**). Strikingly, mice treated with the combination of carboplatin and m214-sEVs showed disease regression with tumor burden significantly less than the other treated groups by the RW1 timepoint (**Fig. 3B, C**). More importantly, the combination of carboplatin and m214-sEVs significantly improved overall survival, with all animals surviving beyond 200 days post-recurrence (**Fig. 3D**). These results demonstrate that m214-sEVs significantly enhance the efficacy of carboplatin, providing sustained therapeutic benefit.

**Figure 3.**
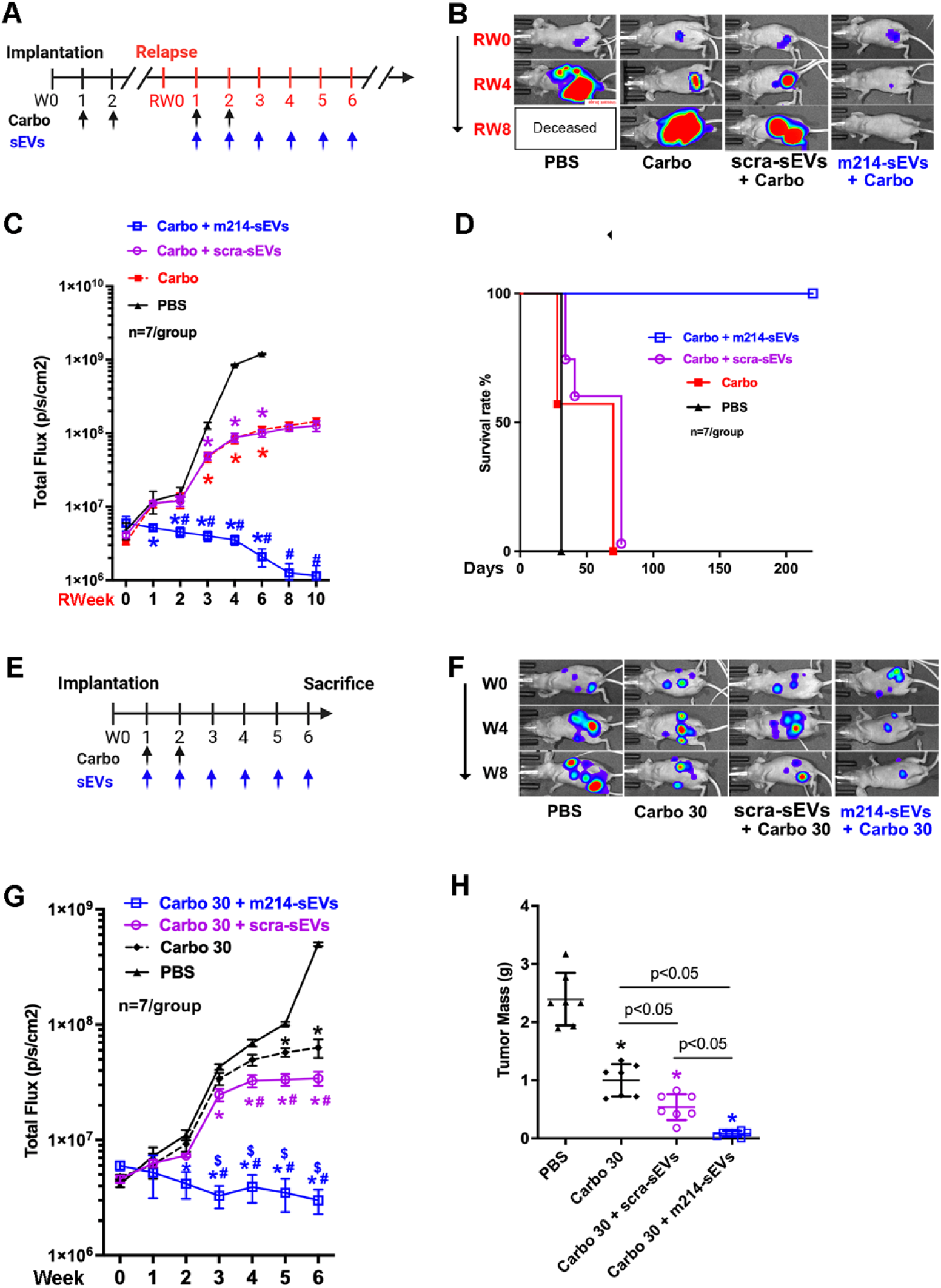
m214-sEVs Enhance Carboplatin Therapeutic Efficacy and Prolong Survival in OC Models. (**A**) Schematic of the treatment timeline for the survival study. Mice bearing intraperitoneal OVCAR3/luc tumors were treated with carboplatin (Carbo, 60 mg/kg, black arrows), and/or sEVs (scra-sEVs or m214-sEVs, blue arrows) beginning at the time of tumor recurrence (RW1). (**B**) Representative BLI of mice show tumor burden across the treatment groups. (**C**) Quantification of BLI signal over time demonstrates reduced tumor progression in the Carbo + m214-sEV group. *p < 0.05 vs. PBS; #p < 0.05 vs. Carbo alone, assessed by two-way ANOVA with Tukey’s post hoc test. (**D**) Kaplan–Meier survival analysis reveals significantly prolonged survival in mice treated with Carbo + m214-sEVs compared to other groups. Log-rank (Mantel–Cox) test results: p < 0.05 for Carbo vs. PBS and Carbo + scra-sEVs vs. PBS; p < 0.001 for Carbo + m214-sEVs vs. Carbo and Carbo + m214-sEVs vs. PBS; non-significant for Carbo vs. Carbo + scra-sEVs. (**E**) Schematic of the treatment timeline of mice bearing intraperitoneal OVCAR3/luc tumors and received low-dose carboplatin (Carbo, 30 mg/kg; black arrows) and/or sEVs (scra-sEVs or m214-sEVs; blue arrows), starting one week post-implantation (W1). Mice were sacrificed at week 6 (W6) for endpoint analysis. (**F**) Representative BLI of mice from each treatment group. (**G**) Quantification of BLI signal over time showed differences in tumor progression among groups. *p < 0.05 vs. PBS; #p < 0.05 vs. Carbo 30 alone; $p < 0.05 vs. Carbo + scra-sEVs; assessed by two-way ANOVA with Tukey’s post hoc test. (**H**) Endpoint quantification of tumor mass at W6. *p < 0.05 vs. PBS; determined by one-way ANOVA with multiple comparisons.

Although carboplatin-based chemotherapy is effective for treating primary OC[9], its dose-dependent toxicities limit long-term use[33]. To test whether m214-sEVs allow chemotherapy dose reduction, mice bearing primary OVCAR3/luc xenografts were treated with low-dose carboplatin (30 mg/kg) plus m214-sEVs or scra-sEVs (**Fig. 3E**). While carboplatin alone reduced tumor growth, addition of scra-sEVs provided modest benefit. In contrast, combining m214-sEVs with low-dose carboplatin produced the greatest tumor suppression (∼45% improvement at week 6; **Fig. 3F-H**). Together, these findings show that engineered m214-sEVs deliver tumor-suppressive miRNAs that potentiate the antitumor activity of carboplatin in both recurrent and primary OC, enabling dose reduction while achieving sustained therapeutic benefit.

Because advanced HGSOC frequently develops resistance to platinum and taxane chemotherapy[34, 35], we evaluated whether m214-sEVs could restore chemosensitivity in resistant OC cells. In cisplatin-resistant A2780cis cells, m214-sEVs markedly reduced the cisplatin IC_50_ at all tested doses (3×10^7^-3×10^9^ particles/mL), whereas scra-sEVs or naïve CEC-sEVs had no effect (**Supplemental Fig. 4A**). Comparable efficacy at 3×10^8^ particles/mL established this as the optimal in vitro dose. In patient-derived paclitaxel-resistant OCSCs (R182 and R2615) [5, 6], m214-sEVs, but not scra-sEVs, significantly enhanced paclitaxel-induced cytotoxicity without affecting cell viability alone (**Supplemental Fig. 4BC**). Together, these results show that m214-sEVs effectively sensitize chemoresistant OC and OCSC models to both platinum and taxane chemotherapy, supporting their use as a therapeutic adjunct for treatment-resistant ovarian cancer.

### m214-sEVs Elevate miR-214-3p/miR-199a-5p and Reduce TLR4 and **β**-catenin in OC

To determine how m214-sEVs sensitize OC to chemotherapy, we first examined their tumor uptake in vivo. Mice bearing recurrent OVCAR3/luc tumor at week 4 post-relapse (**Fig. 4A**) were injected i.p. with GFP-labeled m214-sEVs (GFP-m214-sEVs), which were readily detected within tumor tissues by confocal microscopy and immunogold EM, confirming internalization by both tumor cells and adjacent fibroblasts (**Fig. 4B-E**). GFP signal was not detectable in tumors from mice treated with unlabeled m214-sEVs, confirming the specificity of the GFP signal (**Fig. 4B**). In mice bearing mCherry-labeled OCSC1-F2 tumors, biodistribution imaging using Gaussia luciferase-tagged m214-sEVs (Gluc-m214-sEVs) further revealed selective accumulation in mCherry⁺ tumor nodules with minimal uptake in other organs (**Fig. 4F-G**). In vitro, blocking clathrin-mediated endocytosis with chlorpromazine at the minimal cytotoxic dose (5 µg/mL)[36] abolished the sensitizing effect of m214-sEVs, indicating that clathrin-dependent uptake is required for their function (**Fig. 4H**). These results show that m214-sEVs are internalized by OC tumors and that this is required for their function.

**Figure 4.**
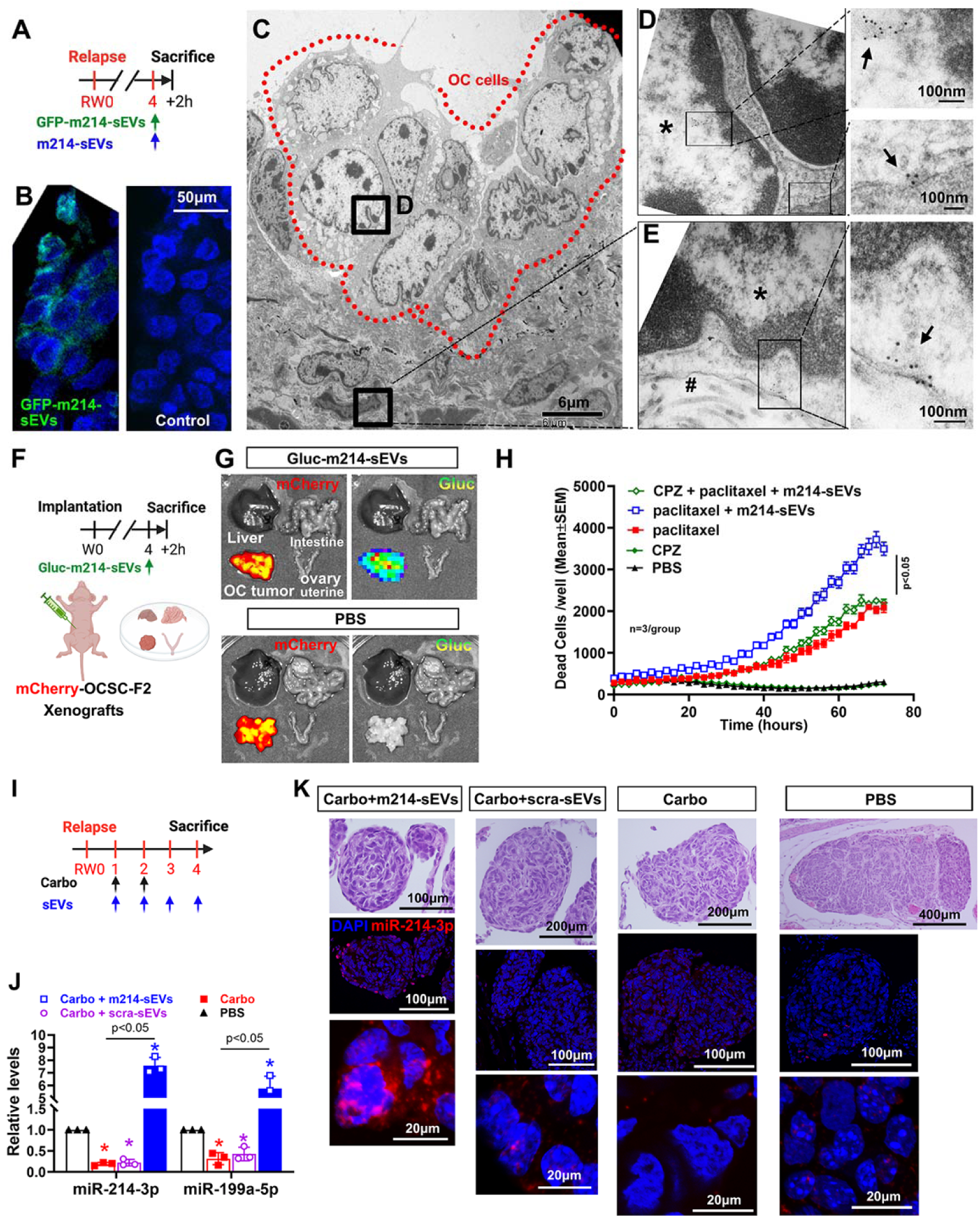
Biodistribution, Cellular Uptake, and Intratumoral miR-214-3p/199a-5p Levels of m214-sEVs Treatment in Recurrent OC. (**A**) Schematic of the experimental timeline for sEV uptake. Mice bearing relapsed OC at relapse week 4 (RW4) were intraperitoneally injected with either GFP-labeled m214-sEVs or unlabeled m214-sEVs and sacrificed 2 hours post-injection. (**B**) Immunofluorescence images of relapsed OC nodules showed strong GFP signal in tumors from mice treated with GFP-m214-sEVs, while no GFP signal was detected in the unlabeled control group. (**C**) Representative TEM image of a relapsed OC nodule, with tumor cells outlined by red dashed lines. (**D**) Higher-magnification view of the boxed region in (C) revealed immunogold-labeled GFP particles localized to both the nucleus (*) and cytoplasm of tumor cells. (**E**) Adjacent stromal region from the same nodule showed immunogold-labeled particles within fibroblast cytoplasm, closely associated with collagen fibers (#). (**F**) Schematic of the experimental design for Gluc-m214-sEV tracking. Mice bearing mCherry⁺ OCSC1-F2 xenografts were intraperitoneally injected with Gluc-labeled m214-sEVs (Gluc-m214-sEVs), and tissues were harvested 2 hours later. (**G**) *Ex vivo* BLI demonstrated preferential accumulation of Gluc-m214-sEVs in OC tumor tissues, with minimal signal detected in liver, ovary, or intestine. (**H**) Real-time CellTox Green cytotoxicity assay demonstrated the effect of pretreatment with chlorpromazine (CPZ) prior to paclitaxel and/or m214-sEVs in OCSC-F182 cells. *p < 0.05, assessed by one-way ANOVA with Tukey’s post hoc test; n = 3 wells/group. (**I**) Schematic of the treatment timeline for miRNA analysis. Mice bearing relapsed OC were treated with carboplatin (Carbo, black arrows) and/or small extracellular vesicles (sEVs, blue arrows), and tumors were collected at relapse week 4 (RW4). (**J**) qRT-PCR analysis of tumor tissues showed significantly elevated levels of miR-214-3p and miR-199a-5p in the combination treatment group (m214-sEVs + Carbo) compared to all other groups. *p < 0.05, one-way ANOVA with Tukey’s post hoc test. (**K**) Representative H&E staining and fluorescent in situ hybridization (ISH) images of relapsed OC nodules revealed increased miR-214-3p expression (red signal) in OC cells following combination treatment with m214-sEVs and carboplatin.

We next investigated whether m214-sEVs modulate miR-214-3p and miR-199a-5p levels in recurrent OC tumors. qRT-PCR of recurrent OVCAR3/luc tumors collected at week 4 post-treatment (**Fig. 4I**) showed that carboplatin + m214-sEVs significantly increased miR-214-3p and miR-199a-5p expression compared with carboplatin alone or with scra-sEVs (**Fig. 4J**). Fluorescent ISH (FISH) confirmed restoration of miR-214-3p levels within tumor tissues (**Fig. 4K**). These findings demonstrate that m214-sEVs restore and elevate chemotherapy-suppressed miR-214-3p/miR-199a-5p expression in the TME

TLR4 and β-catenin are validated targets of miR-214-3p [37, 38] and key mediators of chemoresistance [15, 16, 39]. Our previous studies demonstrated that miR-199a-5p regulates chemoresistance through the TLR4 pathway[15] and that β-catenin is selectively upregulated in chemoresistant OCSCs[39]. We next analyzed their expression in recurrent OC. Western blot analysis of recurrent OC tumors isolated from mice carrying recurrent OVCAR3/luc tumor after 4 weeks of treatment (**Fig. 5A**) showed that carboplatin alone elevated both proteins, whereas the combination with m214-sEVs markedly reduced their levels and increased cleaved caspase-3, indicating enhanced apoptosis (**Fig. 5BC**). Similarly, in paclitaxel-resistant OCSC-R182 cells, m214-sEVs suppressed paclitaxel (20 µM)-induced upregulation of TLR4 and β-catenin while enhancing caspase-3/7 activity (**Fig. 5D-F**). To confirm that the therapeutic effect depends on encapsulated vesicle cargo, protease protection assay showed that proteinase K or RNase treatment in the presence of detergent Triton X-100 (which disrupts vesicle membranes and abolishes sEV protection) [40] abolished m214-sEV cytotoxicity, whereas enzyme treatment alone did not (**Supplemental Fig. 5**), demonstrating that encapsulated miRNAs, not surface molecules, mediate the effect.

**Figure 5.**
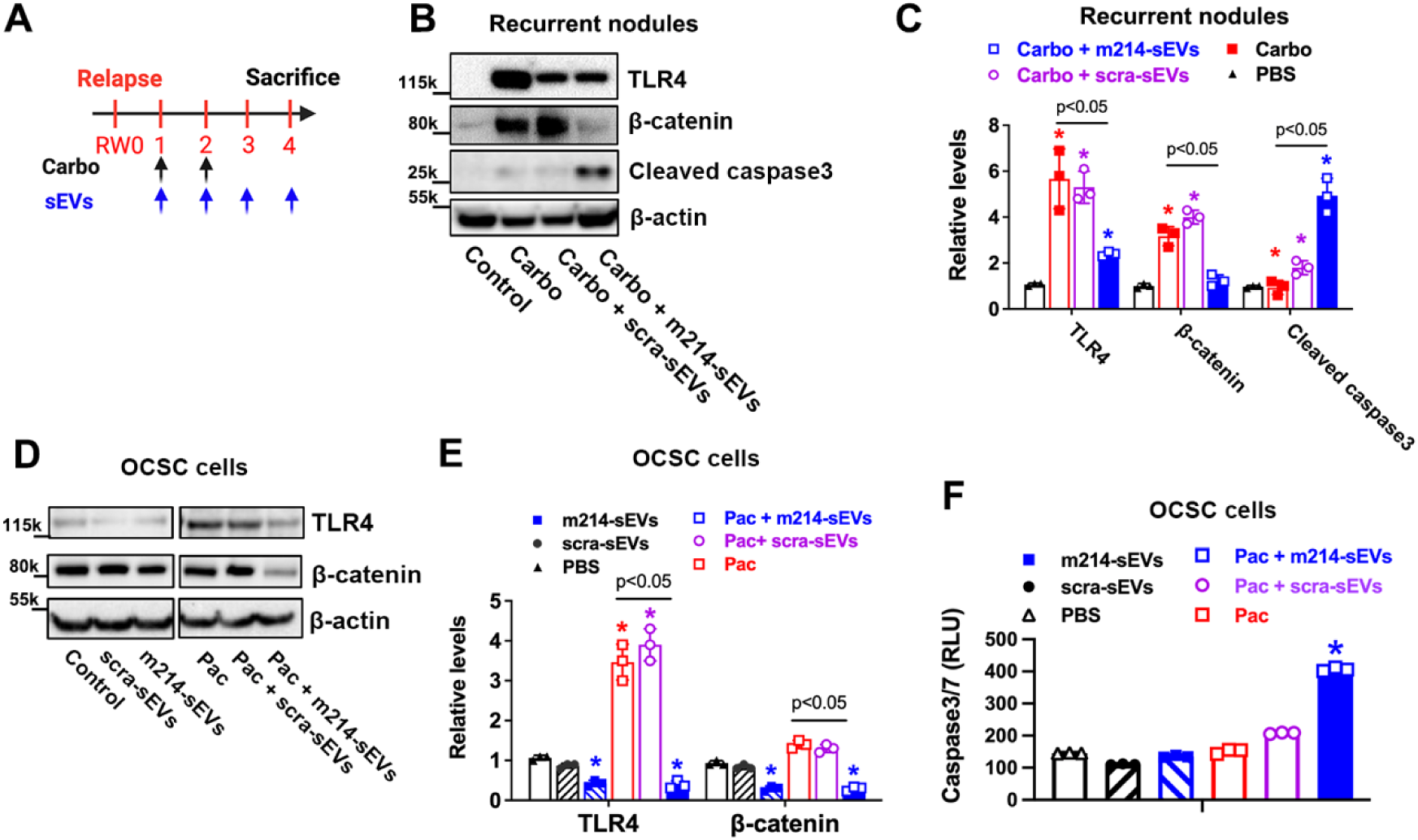
Combined m214-sEV and Chemotherapy Treatment Downregulates TLR4/β-Catenin and Promotes Apoptosis in OC. (**A**) Schematic of the treatment schedule. Mice bearing relapsed OC tumors were treated with carboplatin (black arrows) and/or sEVs (blue arrows). Tumors were harvested four weeks after recurrence. (**B**) Representative Western blots and (**C**) corresponding quantitative analysis of tumor lysates showed reduced expression of TLR4 and β-catenin, and increased levels of cleaved caspase-3 in mice receiving the combination treatment with m214-sEVs and carboplatin. *p < 0.05, assessed by one-way ANOVA with Tukey’s post hoc test. (**D, E**) Western blot (**D**) and quantitative analysis (**E**) of TLR4 and β-catenin expression in OCSC-R182 cells following indicated treatments. (**F**) Caspase-3/7 activity assay in OCSC1-F2 cells showed increased apoptosis following combined treatment with m214-sEVs and paclitaxel (Pac). *p < 0.05 vs. Pac or other treatment groups, assessed by one-way ANOVA.

Together, these data show that m214-sEVs deliver functional miR-214-3p/miR-199a-5p to recurrent OC, downregulate their targets TLR4 and β-catenin, and enhance apoptosis and chemosensitivity to both platinum and taxane therapies.

### m214-sEV and Carboplatin Combination Therapy Reprograms Secondary Tumor-Derived sEVs Toward a Less Pro-Tumorigenic Phenotype

Further ultrastructural analysis of immunogold-labeled recurrent OVCAR3/luc tumors from mice at four weeks post-relapse (**Fig. 6A**), treated with GFP-m214-sEVs in combination with carboplatin revealed abundant GFP-positive signals within intraluminal vesicles of multivesicular bodies (MVBs) in OC cells (**Fig. 6B**). These findings indicate the presence of internalized m214-sEVs within the MVB compartment, suggesting the involvement of sEV biogenesis in treated OC cells. GFP signals were also detected on vesicular structures in the adjacent extracellular space (**Fig. 6C**), implying that m214-sEVs may interact with neighboring tumor niche cells. Primary tumor cell-derived sEVs (t-sEVs) are known to promote malignancy [41, 42]; however, emerging evidence[43, 44], including the concept proposed by Askenase[27], suggests that secondary EVs released by the tumor and its niche cells that internalize exogenous EVs can substantially shape the overall biological outcomes. To test this, we harvested tumor tissues from recurrent OC-bearing mice four weeks after the initiation of the combination treatment and cultured them *ex vivo* for 24 hours to collect secondary t-sEVs (**Fig. 6D**). Using these t-sEVs (3×10L particles/mL), we then treated mCherry-labeled chemoresistant OCSC1-F2 cells[6]. Secondary t-sEVs from PBS- or carboplatin-only–treated mice (PBS-t-sEVs and carbo-t-sEVs) significantly increased the IC₅₀ of carboplatin, suggesting that these t-sEVs promote chemoresistance. In contrast, t-sEVs isolated from mice that received the combination treatment (m214 + carbo-t-sEVs) significantly reduced the cisplatin IC₅₀, indicating a loss of their pro-resistance function (**Fig. 6E**). Western blot analysis revealed that Carbo-t-sEVs contained elevated levels of integrin β1 and matrix metalloproteinase-9 (MMP9), both of which are associated with extracellular matrix remodeling, stemness, invasion, and chemoresistance[45, 46]. In contrast, m214+Carbo-t-sEVs showed significantly reduced levels of integrin β1 and MMP9 compared to both PBS-t-sEVs and Carbo-t-sEVs (**Fig. 6F-G**). Interestingly, MMP2 levels remained unchanged across all treatment groups. Consistent with the protein data, invasion assays showed that PBS-t-sEVs and Carbo-t-sEVs enhanced the invasive capacity of OCSC-F2 cells, whereas m214+Carbo-t-sEVs had no such effect, indicating a loss of pro-metastatic potential (**Fig. 6H**). Collectively, our findings suggest that secondary t-sEVs generated from m214-sEV-recipient tumor and niche cells may transition from a pro-tumorigenic to a less tumor-promoting or even an anti-tumorigenic phenotype, thereby amplifying the therapeutic impact of m214-sEVs.

**Figure 6.**
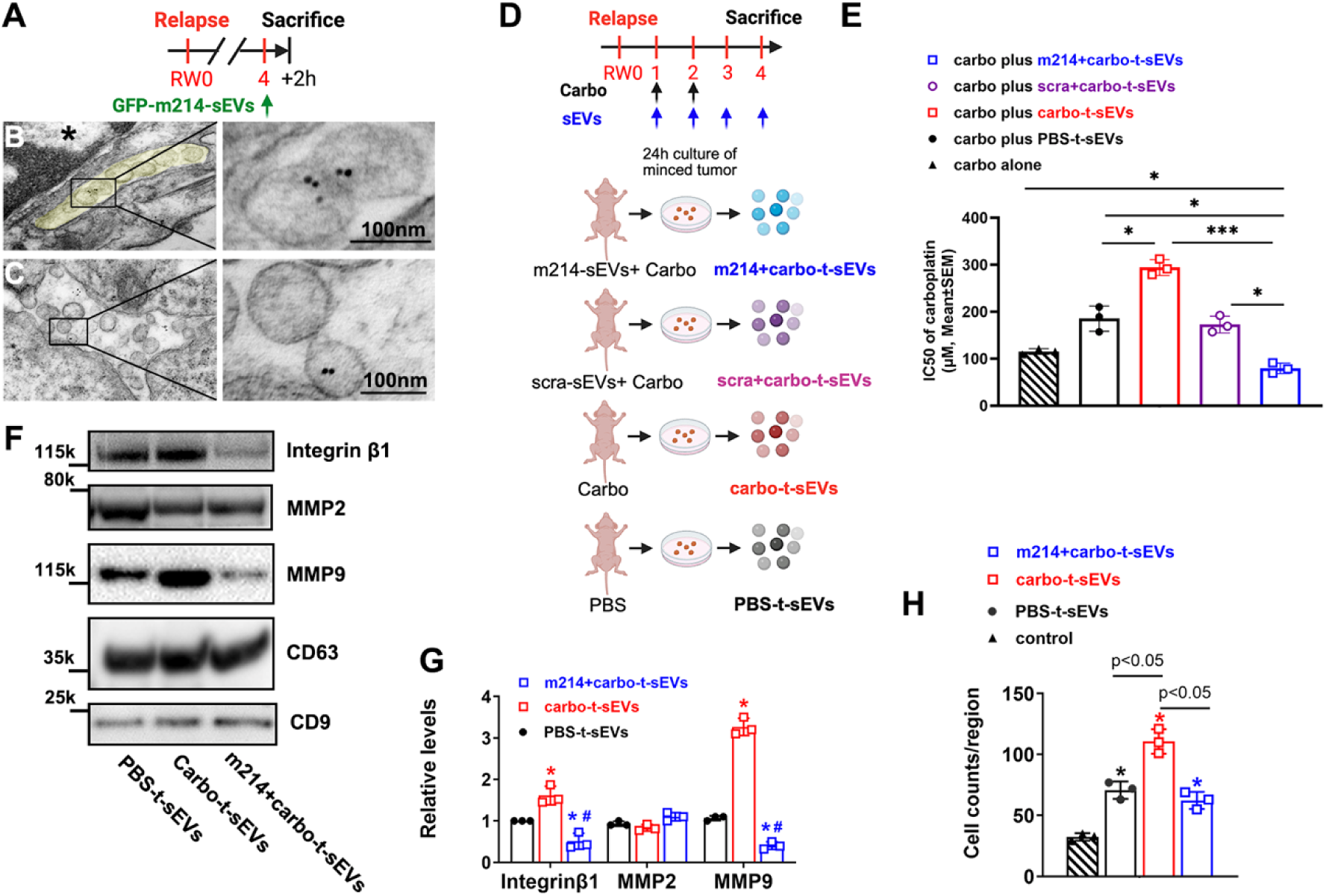
m214-sEVs Change Secondary t-sEV Composition and Abrogate Its Pro-metastatic Effects in OC. (**A**) Schematic illustrating the in vivo treatment regimen and timeline for tissue collection. (**B, C**) Representative TEM images of relapsed OC tissue showed immunogold-labeled GFP-m214-sEVs localized within MVBs (**B**) and in the extracellular space (**C**). (**D**) Schematic of experimental design: tumor-derived sEVs (t-sEVs) were isolated from minced tumor tissues of mice treated under four different conditions. (**E**) Quantification of carboplatin IC₅₀ values in mCherry⁺ OCSC-F2 cells following treatment with various t-sEVs preparations. (**F, G**) Western blot analysis (**F**) and corresponding quantification (**G**) of t-sEV cargo proteins, including Integrin β1, MMP2, MMP9, CD63, and CD9. (**H**) Quantitative results of transwell invasion assays in OCSC-F2 cells treated with t-sEV. *p < 0.05, **p < 0.01, ***p < 0.001 by one-way ANOVA with Tukey’s post hoc test.

### YKT6 Downregulation Mediates the Anti-Migratory and Anti-Tumor Effects of m214-sEV and Carboplatin Combination Therapy in Recurrent Ovarian Cancer

We found that treatment with m214-sEVs in combination with carboplatin significantly reduced the expression of YKT6, a high-confidence predicted target of miR-214-3p identified by TargetScan [47](weighted context++ score = −0.06) (**Fig. 7A**), in ovarian tumors isolated from mice bearing recurrent OVCAR3/luc xenografts after four weeks of treatment (**Fig. 7BC**). Consistently, proteomic profiling of OVCAR3 cells treated with m214-sEVs revealed a downregulation of YKT6 (**Supplemental Excel 1**). YKT6 is a member of the vesicular soluble N-ethylmaleimide-sensitive factor attachment protein receptor (v-SNARE) family [48], and promotes metastasis in lung and breast cancers by driving epithelial–mesenchymal transition (EMT) and enhancing MMP activity[49, 50].

**Figure 7.**
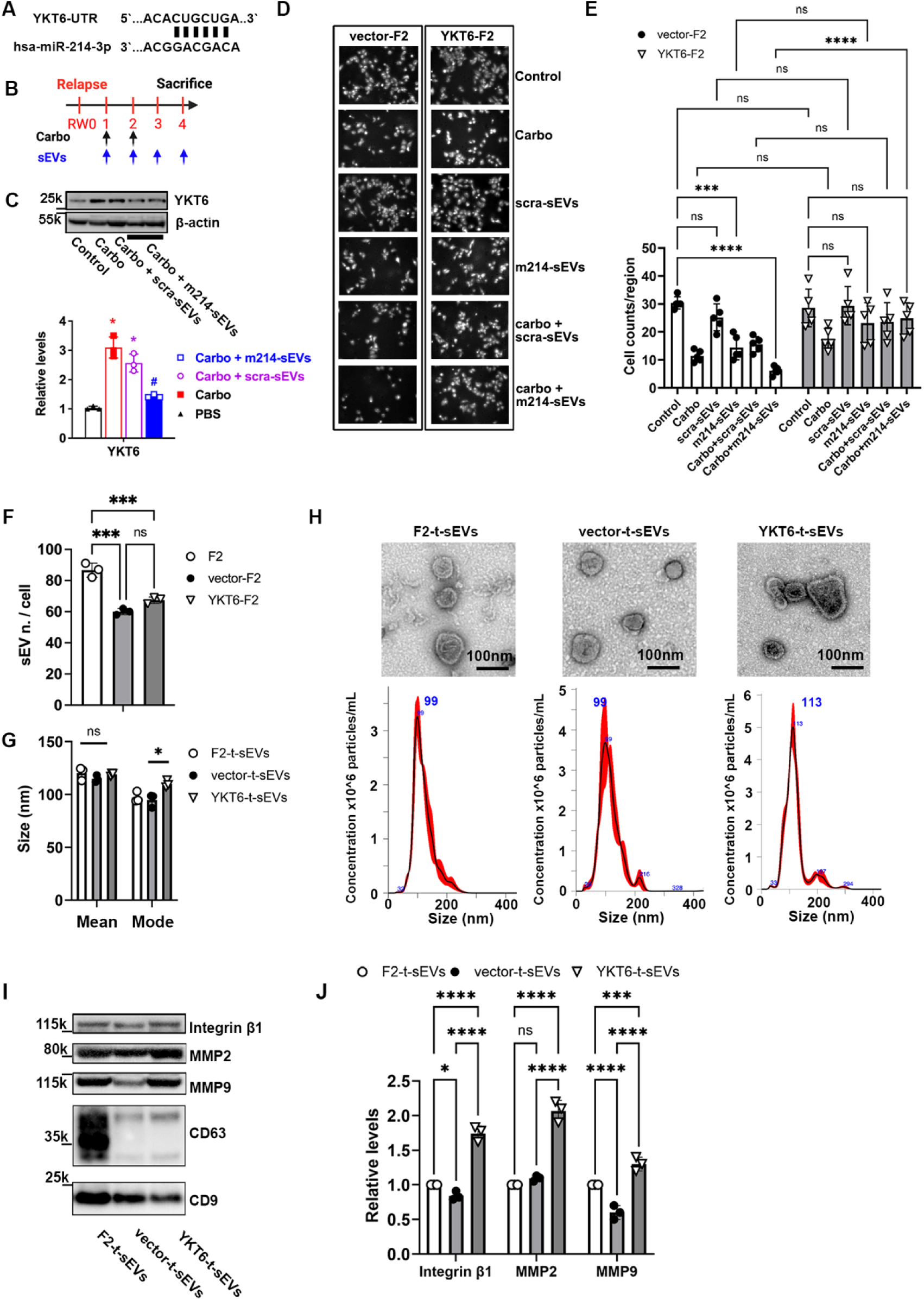
Functional Role of YKT6 in Cell Motility and sEV Cargo in OC. (**A**) Schematic of the treatment timeline showing administration of carboplatin (Carbo) and/or sEV, followed by tissue collection at relapse week 4 (RW4). (**B**) Representative Western blot and corresponding quantification of YKT6 protein levels in recurrent OC xenografts. (**C**) Schematic illustrating the predicted miR-214-3p binding site within the 3’UTR of YKT6 mRNA. (**D**) Representative images and quantification of transwell migration assays comparing YKT6-overexpressing (YKT6-F2) and control vector-transduced (vector-F2) cells. (**E**) Quantification of sEV secretion per cell, measured by NTA, in YKT6-F2, vector-F2, and F2 parental cells following the indicated treatments. (**F**) Quantification of mean and mode sizes of t-sEVs from different cell types. (**G**) Representative TEM images and corresponding NTA profiles showing t-sEV morphology and size distribution. (**H**) Representative Western blot and (**I**) quantification of protein cargo in three types of t-sEVs. *p < 0.05, ***p < 0.001, ****p < 0.0001, ns = not significant; assessed by one-way ANOVA with Tukey’s post hoc test.

To test its role in therapy response, we generated YKT6-overexpressing OCSC1-F2 cells (YKT6-F2) via retroviral transduction (Addgene #116946, pMRXIP-GFP-YKT6). Western blot analysis confirmed successful YKT6 overexpression in YKT6-F2 cells compared to control vector-transduced F2 cells (vector-F2) (**Supplemental Fig. 6 AB**). Migration assays showed that treatment of vector-F2 cells with the combination of m214-sEVs and carboplatin significantly suppressed cell migration compared to either scra-sEVs plus carboplatin or carboplatin alone (**Fig. 7DE**). In contrast, this combination treatment did not reduce the migration of YKT6-F2 cells, indicating that YKT6 overexpression abrogates the anti-migratory effect (**Fig. 7DE**). Notably, no significant differences in migration were observed between vector-F2 and YKT6-F2 cells under PBS, monotherapies (carboplatin, scra-sEVs, m214-sEVs), or the scra-sEV plus carboplatin treatment (**Fig. 7DE**). These findings suggest that the inhibitory effect of m214-sEV and carboplatin combination treatment on F2 cell migration is specific and YKT6-dependent. Together, these *in vitro* results support a role for YKT6 downregulation in mediating the therapeutic activity of m214–sEV–based combination treatment of recurrent OC.

Given YKT6’s established role in EV biogenesis and cargo loading[51, 52], we next examined whether YKT6 overexpression influences t-sEV production and content. NTA and TEM results revealed no significant differences in particle concentration or morphology between YKT6-F2, vector-F2, and parental OCSC1-F2 cells (**Fig. 7F-H, Supplemental Fig. 6C**). However, sEVs from YKT6-F2 cells (YKT6-t-sEVs) exhibited significantly increased particle size (**Fig. 7G**) and were enriched with pro-metastatic proteins, including integrin β1, MMP2, and MMP9, compared to control vector-derived sEVs (**Fig. 7IJ**). Because YKT6 is a SNARE protein that regulates vesicular trafficking pathways influencing key cellular processes such as adhesion[53, 54], our *in vitro* findings, together with the observed *in vivo* downregulation of YKT6 by the m214-sEV and carboplatin combination therapy, support a model in which YKT6 suppression enhances the therapeutic efficacy of the combination treatment by limiting the production of pro-tumorigenic t-sEVs in recurrent OC.

..

## Discussion

Recurrent OC remains incurable, primarily due to the development of chemoresistance and metastatic progression[3, 4]. Clinically, patients with advanced-stage OC frequently exhibit marked downregulation of the miR-214-3p/miR-199a-5p cluster, and higher expression levels of this cluster are correlated with improved tumor-free survival (**Fig. 1**). In this study, we demonstrate that engineered m214-sEVs, enriched with miR-214-3p and miR-199a-5p, restore chemosensitivity and suppress tumor growth in pre-clinical models of recurrent OC. Mechanistically, these therapeutic effects are mediated, at least in part, through the coordinated downregulation of chemoresistance-associated genes such as TLR4 and β-catenin, and through inhibition of the SNARE protein YKT6 (**Fig. 8**). Our findings establish a therapeutic paradigm in which engineered sEVs deliver clinically lost tumor-suppressive miRNAs to reprogram chemoresistance-associated gene networks and remodel the TME in recurrent OC.

**Fig. 8.**
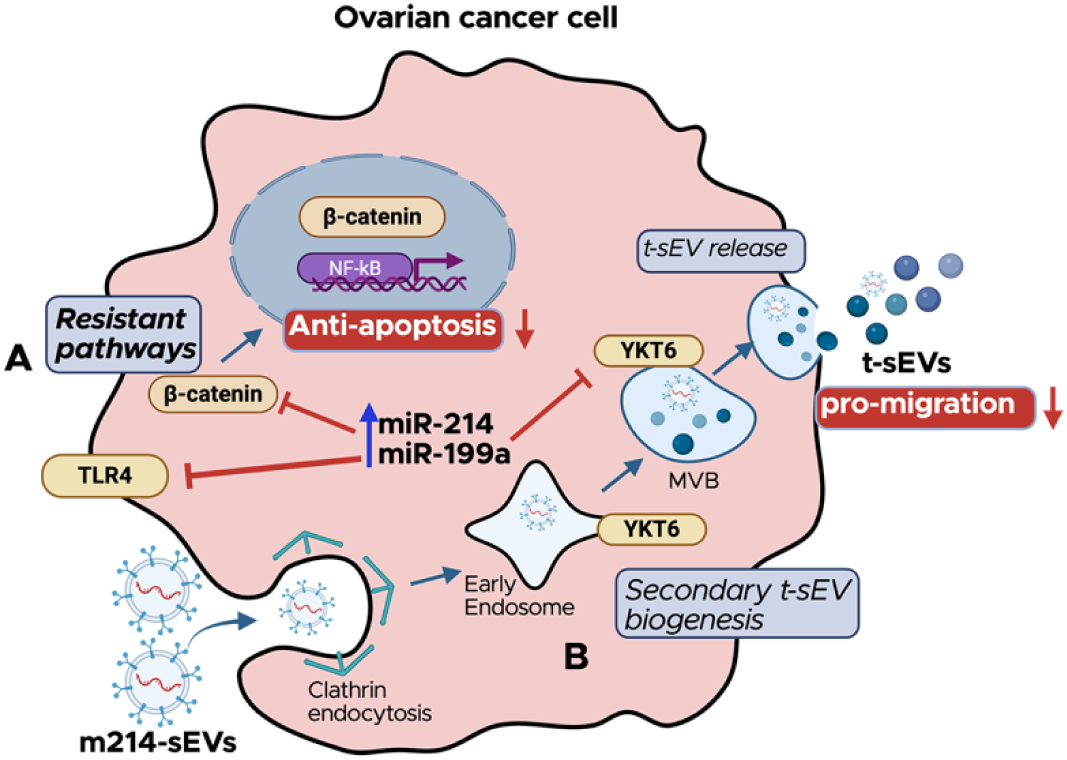
Schematic Illustrating the Proposed Molecular and Cellular Mechanisms by which m214-sEVs Exert Therapeutic Effects in Recurrent OC. **(A)** Following clathrin-mediated endocytosis, m214-sEVs deliver miR-214-3p and miR-199a-5p into ovarian cancer cells, resulting in the downregulation of chemoresistance-associated genes, TLR4 and β-catenin. This suppression inhibits anti-apoptotic signaling pathways and restores sensitivity to chemotherapy. (**B**) m214-sEVs also downregulate YKT6, a SNARE protein involved in MVB trafficking, leading to altered biogenesis and protein cargo composition of secondary tumor-derived sEVs (t-sEVs), contributing to reduced tumor invasiveness and recurrence. Created in BioRender (BioRender.com/2fgsdlc).

We employed a clinically relevant OC mouse model that recapitulates relapse following first-line platinum-based chemotherapy (**Fig. 1**). Unlike models that rely on prolonged drug exposure or artificial induction of resistance, this model enables spontaneous OC recurrence, capturing the complexity of intratumoral heterogeneity, including OCSCs, stromal fibroblasts, and immune cell populations[6, 55, 56]. Importantly, this model mirrors the stage-dependent dysregulation of miR-214-3p/miR-199a-5p observed in patients, in which downregulation of these tumor-suppressive miRNAs parallels disease progression. In situ hybridization revealed consistent downregulation of the miR-214-3p/miR-199a-5p cluster in recurrent OC tumors and human HGSOC, while adjacent stromal cells retained expression of this cluster, highlighting a cell-type–specific and progression-linked pattern of miRNA loss. Treatment with m214-sEVs in combination with carboplatin resulted in sustained tumor suppression for more than 200 days, whereas monotherapies did not block tumor recurrence (**Fig. 3**). In vitro, m214-sEVs enhanced chemosensitivity across multiple chemoresistant and recurrent OC cell lines. Moreover, internalization of m214-sEVs by tumor cells increased intracellular levels of the miR-214-3p/miR-199a-5p cluster (**Fig. 4**). Together, these findings demonstrate that miRNA dysregulation in OC is both stage-specific and context-dependent, and that targeted restoration of clinically lost tumor-suppressive miRNAs via m214-sEVs can effectively overcome chemoresistance and prevent tumor recurrence.

sEVs are increasingly recognized as promising therapeutic vehicles due to their biocompatibility and ability to deliver functional cargo to recipient cells[19, 20]. However, the intracellular mechanisms by which EV cargo exerts its effects on recipient cells remain incompletely understood. We selected CEC-sEVs based on our previous findings that these vesicles enhance chemotherapy efficacy. In this study, m214-sEVs, but not scra-sEVs, elevated miR-214-3p/miR-199a-5p levels in tumor tissues, indicating that selective enrichment of therapeutic miRNA cargo is required to achieve targeted biological effects. Notably, m214-sEVs suppressed the expression of known miR-214-3p/miR-199a-5p target genes, TLR4, β-catenin, and YKT6, which are key regulators of tumor progression, metastasis, and chemoresistance (**Fig. 5**). A gain-of-function experiment revealed that YKT6 overexpression in OCSCs abrogated the anti-migratory effects of the m214-sEV and carboplatin combination treatment, implicating YKT6 as a critical functional target of the therapy (**Fig. 7**). Moreover, suppression of TLR4 and β-catenin sensitized OCSCs to paclitaxel (**Fig. 5**), although the precise mechanistic link between individual miRNA targets and responsiveness to different chemotherapeutic agents remains to be defined.

Compared to primary tumor-derived sEVs (t-sEVs), the contribution of secondary t-sEVs in modulating tumor progression remains understudied [27]. The present study showed that secondary tEVs, generated by tumor and niche cells that internalized m214-sEVs following combination treatment with m214-sEVs and carboplatin, convert primary t-sEVs from a pro-tumorigenic to an anti-tumorigenic state, as evidenced by a marked reduction in cargo proteins such as integrin β1 and MMP9 (**Fig. 6**). Consistent with the established role of YKT6 in endosomal membrane cycling and cargo sorting [51, 52], our data suggest that YKT6 plays a key role in determining the protein composition of t-sEVs. Specifically, t-sEVs derived from YKT6-overexpressing OCSCs were enriched in pro-metastatic proteins and could override the anti-migratory effects of m214-sEV and carboplatin combination treatment (**Fig. 7**). Thus, YKT6 suppression not only limits tumor invasion, but also curtails the release of pro-metastatic secondary t-sEVs that can shape the TME, extending the therapeutic benefit of m214-sEVs beyond direct tumor cell targeting. This supports the emerging concept proposed by Askenase [27] that secondary EVs, in concert with m214-sEVs, collectively shape the biological outcome of the combination intervention.

Furthermore, m214-sEVs were taken up by adjacent stromal fibroblasts, further indicating that the combination treatment may also influence the TME by modulating non-malignant niche cells (**Fig. 4**). Given that secondary EV-mediated signaling plays a critical role in regulating immune cell responses, particularly macrophage polarization [27, 43, 44], we speculate that the therapeutic efficacy of m214-sEVs in combination with carboplatin may, in part, be driven by immunomodulatory mechanisms. This potential effect on the immune compartment warrants further investigation to fully elucidate the role of EV-mediated crosstalk in the anti-tumor activity of the combination therapy.

Together, these findings provide compelling evidence that m214-sEVs, when combined with carboplatin, represent a promising therapeutic strategy to overcome chemoresistance and prevent recurrence in OC. By delivering tumor-suppressive miRNAs and modulating both tumor-intrinsic pathways and the composition of secondary tumor-derived sEVs, this combination therapy targets multiple mechanisms of disease progression. Importantly, the ability of m214-sEVs to alter the TME, including potential effects on immune modulation, further enhances their therapeutic relevance. Given their biocompatibility, targeted delivery capabilities, and scalable production, engineered sEVs represent a clinically translatable platform [57, 58] that may improve outcomes for patients with recurrent OC, a population with limited treatment options. The present study provides proof-of-concept that precision restoration of clinically depleted miRNA networks through engineered sEVs offers a next-generation therapeutic modality for recurrent ovarian cancer.

This study has several limitations. The m214-sEV dose was selected based on prior data[9, 30]; future work will define dose-response relationships and optimal therapeutic windows. While ultrastructural data suggest m214-sEV recycling through MVBs, distinguishing between secondary tEVs generated via MVB recycling and those released directly from the cytosol requires further investigation. Such studies will also help determine whether miR-214/miR-199a are incorporated into secondary EVs to mediate downstream biological effects.

## Materials and methods

### Human Subjects

Human ovarian cancer samples were collected with approval from the Wayne State University Institutional Review Board (IRB-20-07-2521) and the Karmanos Cancer Institute Institutional Review Board (IRB-2013-052). All samples were obtained following informed consent and subsequently de-identified. Research involving human subjects was conducted in full compliance with institutional guidelines and actively reviewed by the respective IRBs.

### TCGA Data Analysis

Expression profiles of hsa-miR-214-3p (MIMAT0000271) and hsa-miR-199a-5p (MIMAT0000231) were obtained from TCGA ovarian serous cystadenocarcinoma dataset (TCGA-OV) via the Genomic Data Commons portal (https://portal.gdc.cancer.gov)[59]. miRNA-seq data generated by the British Columbia Genome Sciences Centre (BCGSC) pipeline were normalized as reads per million (RPM) and log₂-transformed (log₂[value + 1]). Corresponding clinical data were downloaded from the same cohort, excluding samples without clinical information.

For clinical correlation analysis, patients were stratified by histological grade (G1+G2 vs G3+G4), and differences in miRNA expression were compared using the Wilcoxon rank-sum test. Statistical analyses were performed using R (v4.2.1) with the stats, car, deseq2, and ggplot2 packages [60].

For prognostic evaluation in the tumor-free patient subset who were annotated as having no residual disease or recurrence at last follow-up, survival analyses were performed in R using the survival, survminer, and ggplot2 packages[61]. Optimal expression cut-off points were determined by the surv_cutpoint function to dichotomize patients into high- and low-expression groups. The proportional hazards assumption was tested before model fitting. Kaplan–Meier curves and log-rank tests assessed associations between miRNA expression and overall survival, with P < 0.05 considered statistically significant.

### Cell Lines and Culture Conditions

Human cerebral endothelial cells (CECs) (ACBRI376, Cell Systems) were cultured and passaged in Complete Classic Medium (4Z0-500, Cell Systems). To prepare conditioned medium for sEV isolation, the culture was switched to serum-free medium (SF-4Z0-500, Cell Systems) for 48 hours before collection[9, 26].

The cisplatin-resistant human ovarian cancer (OC) cell line A2780cis (RRID:CVCL_1942, Millipore-Sigma, 93112517) was maintained in RPMI-1640 medium supplemented with 10% fetal bovine serum (FBS). To maintain cisplatin resistance, 1 µM cisplatin was added every two passages[62].

Human ovarian cancer stem cell (OCSC) lines, including R182, R2615, and mCherry-OCSC1-F2, were established as previously described [6] and cultured in RPMI-1640 supplemented with 10% FBS, 1% MEM Non-Essential Amino Acids (ThermoFisher, 11140050), 1% HEPES (ThermoFisher, 15630080), and 1% sodium pyruvate (ThermoFisher, 11360070).

OVCAR3/luc cells were generated by transfecting OVCAR3 cells (RRID: CVCL_0465, ATCC HTB-161) with a luciferase expression vector (pLVX-EF1α-IRES-Puro, Clontech), as previously described[9]. These cells were cultured in RPMI-1640 with 10% FBS and selected with 1.2 µg/mL puromycin.

### Overexpression of MiRNAs in CECs

Lentiviral constructs carrying the human miR-214-3p precursor (PMIR-214-PA-1, System Biosciences) or a scrambled control (PMIRH000PA-1) were used to generate lentivirus using the Lenti-X Packaging System (Takara)[30]. CECs were infected with the viral supernatant, and stably transduced cells were selected using 1.1 µg/mL puromycin. Puromycin-resistant CECs were expanded and used for downstream experiments [26, 30].

### Overexpression of YKT6 in OCSCs

Retroviral constructs carrying GFP-YKT6 (pMRXIP-GFP-YKT6) and the corresponding backbone control vector were used to generate retrovirus using pUMVC (Addgene #8449) and pCMV-VSV-G (Addgene #8454) to co-transfect into HEK293T cells. pMRXIP-GFP-YKT6 was a gift from Noboru Mizushima (Addgene plasmid # 116946 ; http://n2t.net/addgene:116946 ; RRID:Addgene_116946). pUMVC (Addgene plasmid # 8449 ; http://n2t.net/addgene:8449 ; RRID:Addgene_8449) and pCMV-VSV-G (Addgene plasmid # 8454 ; http://n2t.net/addgene:8454 ; RRID:Addgene_8454) were gifts from Bob Weinberg.

OCSCs were infected with the viral supernatant and selected using puromycin at a concentration of 2 µg/mL. Stable, GFP-positive single-cell clones were isolated, expanded, and maintained for downstream experiments.

### The Isolation and Characterization of sEVs

sEVs were isolated from conditioned medium using differential ultracentrifugation, as previously described[7, 9, 17, 30], and were characterized following the MISEV 2018 and 2023 guidelines[7, 9, 17, 30–32]. The concentration and size distribution of CEC-sEVs were analyzed using nanoparticle tracking analysis (NTA) with a NanoSight NS300 (Malvern). The ultrastructural morphology of sEVs was assessed via transmission electron microscopy (TEM; JEM-1400Flash, JEOL) and cryo-electron microscopy (CryoEM; Talos Arctica, ThermoFisher). The expression of sEV surface markers CD63, CD81, and CD9 was evaluated using Western blotting and ExoView analysis, following our previously published protocols[7, 17].

### The Protein Profile of sEVs and Bioinformatics Analysis

Total proteins were extracted from isolated sEVs and analyzed by mass spectrometry-based proteomics, as described in our previous studies[63, 64]. Raw data were processed using Proteome Discoverer 2.4 (Thermo Scientific) and further analyzed with Scaffold software (Proteome Software, Inc.). The raw protein identification results are provided in the Supplemental Excel 1.

Protein enrichment analysis was conducted using the Enrichr gene enrichment analysis tool[65], and Gene Ontology (GO) analysis was performed to identify top-ranked molecular function (MF) categories. First, the enriched proteins are ranked by the spectrum number, and their IDs are converted to the corresponding gene IDs. The gene list was input into the Enrichr website, and the Ontologies database was then applied to perform analysis.

### CEC-sEV Labelling and Tracking

To visualize and track CEC-derived sEVs *in vivo*, CECs were transfected with a plasmid encoding a Gluc-Lactadherin fusion construct (a gift from Dr. Takahashi) [66], which directs Gaussia luciferase (Gluc) expression to sEVs via exosomal lactadherin.

For tracking sEV distribution in OC cells, CECs were also transfected with a plasmid encoding CD63-EGFP (pEGFP-CD63, a gift from Paul Luzio; Addgene plasmid #62964), enabling GFP labeling of sEVs via CD63[9, 17, 30].

Labeled sEVs were administered *in vivo* 2 hours before tissue collection. For detection, immunofluorescent staining was performed on 8 µm cryosections using an anti-GFP antibody (1:500, Cat# 2555, RRID: AB_10692764, Cell Signaling Technology). To determine the subcellular localization of sEVs, immunogold labeling was conducted on 80 nm ultrathin sections using the same primary antibody, followed by 10 nm gold-conjugated streptavidin[9, 17, 30].

### Cell Viability Assays

The cytotoxic effects of sEVs on OC and OCSC cells were assessed using the MTT assay and the CellTox™ Green Cytotoxicity Assay (Promega, G8731), respectively[6, 9].

For the MTT assay, OC cells were seeded in 96-well plates and treated with sEVs and chemotherapy drugs for 72 hours. After treatment, 0.5 mg/mL thiazolyl blue tetrazolium bromide (MTT; MilliporeSigma, M2128) was added to each well and incubated for 4 hours. Absorbance was measured using a microplate reader, and cell viability was calculated relative to untreated controls. The IC₅₀ was determined by linear regression analysis, representing the concentration of treatment required to inhibit 50% of cell viability.

For the CellTox assay, OCSCs were mixed with CellTox Green dye, seeded into 96-well plates, and treated with sEVs and chemotherapy drugs. Fluorescence intensity, corresponding to the number of dead cells, was measured kinetically over 72 hours using a Bio-Rad plate reader.

### Caspase-Glo 3/7 Assay

To assess apoptosis, 10 µg of protein extracted from treated OCSC-R182 cells was diluted in a total volume of 50 µL and mixed with 50 µL of Caspase-Glo 3/7 reagent (Promega). After a 1-hour incubation at room temperature, luminescence was measured using a TD 20/20 Luminometer (Turner Designs, Sunnyvale, CA). Caspase 3/7 activity was calculated relative to untreated controls[5].

### Transwell Migration Assay

OVCAR3 and OCSC1-F2 cells were resuspended in serum-free RPMI-1640 medium and seeded into Transwell inserts (24-well format) at a density of 2.5 × 10L cells in 0.5 mL per insert. The lower chambers were filled with 0.75 mL of complete RPMI-1640 medium containing 10% FBS as a chemoattractant[9].

sEVs and chemotherapy drugs were added to the upper chambers and incubated for 24 hours. Migrated OVCAR3 cells were detected using CellTracker Red CMTPX (ThermoFisher, C34552), while migrated OCSC1-F2 cells were identified by mCherry fluorescence.

### Animals and the *In Vivo* OC Model

All animal procedures were approved by the Henry Ford Hospital Institutional Animal Care and Use Committee (IACUC) under the following protocols: #1634 (April 16, 2018 – April 15, 2021), #1278 (Feb 16, 2021 – Feb 15, 2024), and #1303 (July 14, 2021 – July 13, 2024).

Female BALB/c nude mice (8 weeks old) were used to establish an intraperitoneal OC xenograft model. Mice were injected i.p. with 5×10^6^ OVCAR3/luc cells one week before treatment initiation[9]. Tumor-bearing mice were randomly assigned to receive carboplatin, sEVs, a combination of carboplatin and sEVs, or PBS as a placebo control.

Tumor burden was monitored weekly using BLI with the IVIS Spectrum 200 system (Caliper Life Sciences)[9]. At the endpoint, the mice were sacrificed, the tumors were dissected, and the wet tumor weight was measured.

### In situ Hybridization (ISH) of miRNAs

To detect miRNAs in formalin-fixed, paraffin-embedded (FFPE) human OC tumor sections, the miRNAscope assay (ACD Bio-Techne) was performed according to the manufacturer’s protocol. Pre-designed probes targeting hsa-miR-214-3p (SR-hsa-miR-214-3p-s1), hsa-miR-199a-5p (SR-hsa-miR-199a-5p-s1), a positive control probe for U6 snRNA (SR-RNU6-s1), and a scramble control probe (SR-Scramble-s1) were used. Tissue sections were pretreated with the RNAscope Target Retrieval Kit (322000, ACD Bio-Techne), followed by hydrogen peroxide (H₂O₂) and protease treatment (Pretreat Reagents 322381). Sections were then incubated overnight with the probes. Signal development was performed using the miRNAscope HD Detection Reagents Red (324510). Stained slides were visualized and imaged using a light microscope.

To detect miRNAs in frozen xenograft OC tumor sections from nude mice, FISH was performed based on our published protocols and prior study[67]. Sections were fixed with 1-ethyl-3-(3-dimethylaminopropyl) carbodiimide (EDC) to prevent miRNA degradation. miRCURY LNA™ microRNA detection probes (Exiqon) targeting hsa-miR-214-3p, hsa-miR-199a-5p, U6 snRNA, and a scramble control were used. Fluorescent signals were amplified using TSA PLUS Fluorescence Kits (PerkinElmer), and images were acquired using a confocal microscope.

### The Isolation of sEVs from the OC Tumor

The t-sEVs were isolated according to the published protocol with some modifications[68, 69]. OC xenografts and tumor nodules were dissected from nude mice and rinsed thoroughly with sterile saline to remove blood and debris. Tumor tissues were then minced into small fragments (∼1 mm³) on ice using a sterile razor blade and rinsed again with sterile PBS.

Approximately 0.1 g of tumor tissue was transferred into a 100 mm Petri dish containing 15 mL of OCSC growth medium supplemented with exosome-depleted FBS (Exo-FBS, System Biosciences). After 24 hours of incubation, the supernatant was collected and processed for sEV isolation as described above.

### Quantitative Reverse Transcription-Polymerase Chain Reaction

Total RNA from sEVs and OC xenografts was extracted using the miRNeasy Mini Kit (Qiagen, Cat# 217084). RNA was reverse transcribed using the TaqMan MicroRNA Reverse Transcription Kit (Thermo Fisher Scientific, Cat# 4427975), followed by amplification with TaqMan™ MicroRNA Assays specific for: miR-214-3p (MIMAT0000271, Assay ID: 002306), miR-199a-5p (MIMAT0000231, Assay ID: 000498), miR-15b-5p (MIMAT0000417, Assay ID: 000390), miR-16-2-3p (MIMAT0004518, Assay ID: 002171). U6 snRNA (MIM180692, Assay ID: 001973) was used as the endogenous control. qRT-PCR was performed using the following thermal cycling conditions: 95°C for 20 seconds (initial denaturation), followed by 40 cycles of 95°C for 1 second and 60°C for 20 seconds. Relative miRNA expression levels were calculated using the 2−ΔΔCt method, normalized to U6[9, 30].

### Western Blotting

Proteins were extracted from sEVs, OC cells, or xenograft tumor tissues, and analyzed by Western blotting as previously described in our published protocols[9, 30]. Briefly, 5-10µg proteins were loaded for SDS-PAGE. Primary antibodies were incubated with membranes overnight at 4°C, followed by incubation with secondary antibodies for 2 hours at room temperature (RT). Protein bands were visualized using ECL Western blotting substrates (Thermo Fisher Scientific), and band intensity was quantified using AlphaView SA software (version 3.4.0, Bio-Techne). The original, uncropped images of Western blots are presented in **Supplemental Excel 3**. The following primary antibodies were used:

| Name | Supplier, Catalog # | RRID | Dilution |
| --- | --- | --- | --- |
| CD63 | Santa Cruz, sc-5275 | RRID:AB_627877 | 1:500 |
| CD81 | Abcam, ab109201 | RRID:AB_10866464 | 1:1000 |
| CD9 | Abcam, ab223052 | RRID:AB_2922392 | 1:1000 |
| calnexin | Biolegend, 699041 | - | 1:1000 |
| TLR4 | Novus, NBP2-27149 | RRID:AB_2936296 | 1:1000 |
| β-catenin | Santa Cruz, SC-1496 | RRID:AB_1563968 | 1:1000 |
| Caspase3, cleaved | Cell Signaling, 9661 | RRID:AB_2341188 | 1:1000 |
| MMP2 | MilliporeSigma, MAB13405 | RRID:AB_94120 | 1:1000 |
| MMP9 | Novus, NBP2-41233 | RRID:AB_3302960 | 1:1000 |
| YKT6 | Abclonal, A17083 | RRID:AB_2772927 | 1:1000 |
| Integrin β1 | Santa Cruz, sc-8978 | RRID:AB_2130101 | 1:1000 |
| β-actin | Abcam, ab6276 | RRID:AB_2223210 | 1:5000 |

### Statistical Analysis

All experimental results are based on three independent replicates, unless otherwise indicated with specific n values.

Statistical analyses were performed using GraphPad Prism (version 8.2.1, GraphPad Software). The following tests were applied:

One-way ANOVA with Tukey’s multiple comparisons test for comparisons among more than two groups;

Two-way repeated measures ANOVA for comparisons of OC tumor growth over time between treatment groups;

Unpaired two-tailed Student’s t-test for comparisons between two groups.

Data are presented as mean ± standard error of the mean (SEM). A P-value < 0.05 was considered statistically significant.

## Data Availability Statement

The data that support the findings of this study are available on request from the corresponding authors.

## Acknowledgements

We thank Dr. Paul Stemmer from Wayne State University Proteomics Core for mass spectrometry-based proteomics. The Proteomics Core is supported, in part, by NIH Center grant P30 CA022453 to the Karmanos Cancer Institute at Wayne State University.

