## Supplemental Materials for "Engineered Extracellular Vesicles Enriched with the miR-214/199a Cluster Enhance the Efficacy of Chemotherapy for Ovarian Cancer"

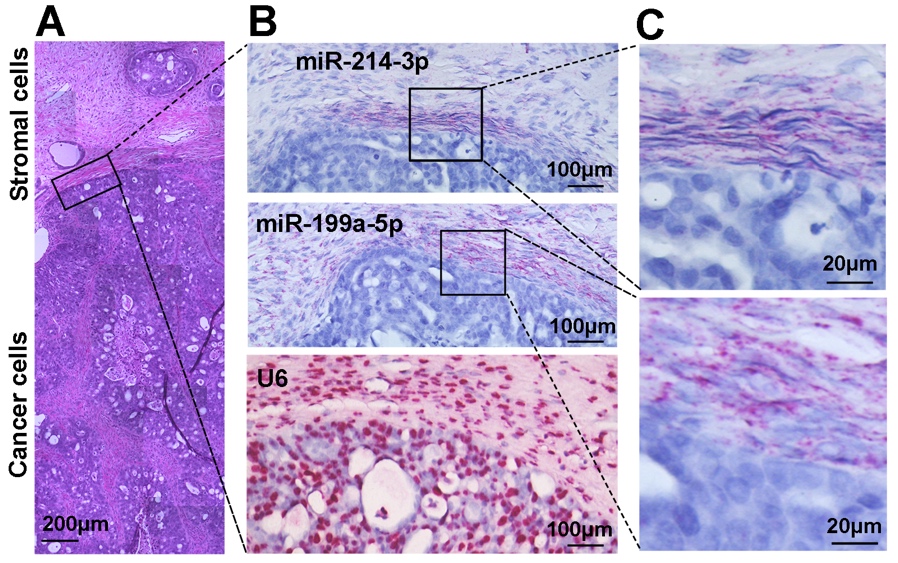

**Supplemental Figure 1. In Situ Detection of miR-214-3p and miR-199a-5p Reveals Stromal Enrichment in Recurrent HGSOC**

(**A**) The H&E-stained section of a representative HGSOC sample of recurrent disease (ID# SKI22475) illustrates demarcated tumor and stromal regions. (**B**) Bright-field images of the selected area from (**A**) following miRNAscope in situ hybridization (ISH) for miR-214-3p and miR-199a-5p, with U6 as a positive control. (**C, D**) Higher-magnification views of the boxed regions in (**B**) showed low expression of miR-214-3p and miR-199a-5p in tumor cells, with stronger signals observed in adjacent stromal compartments.

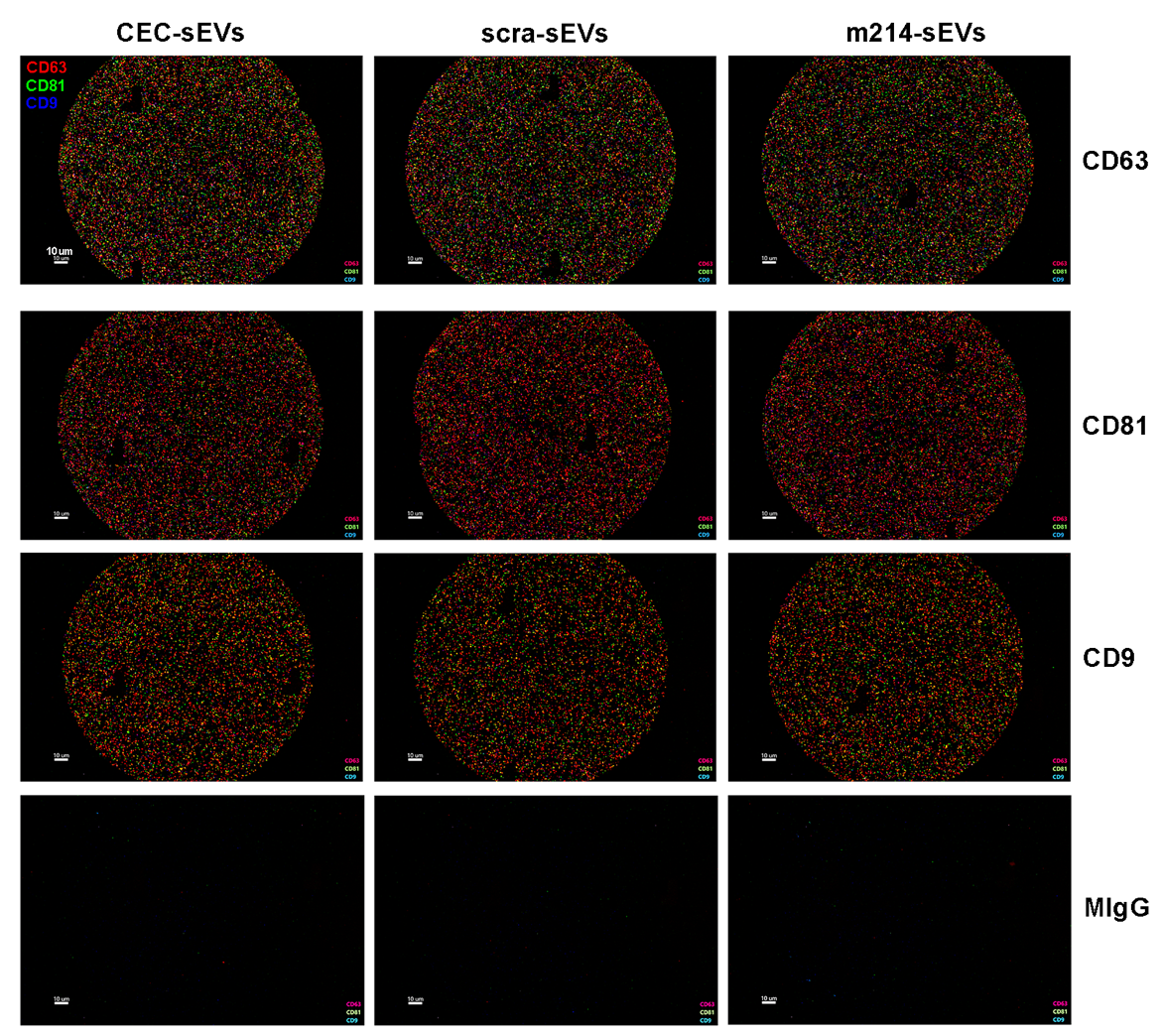

**Supplemental Figure 2. Single EV profiling using the ExoView system**.

Representative fluorescence images acquired using the ExoView R100 platform showed specific capture and labeling of EV particles on chips coated with antibodies against the tetraspanin markers CD63, CD81, and CD9. Strong signal intensities were observed for CEC-sEVs, scra-sEVs, and m214-sEVs on the capture spots. In contrast, minimal fluorescence was detected on control spots coated with mouse IgG (mIgG), confirming the specificity of EV capture and marker detection.

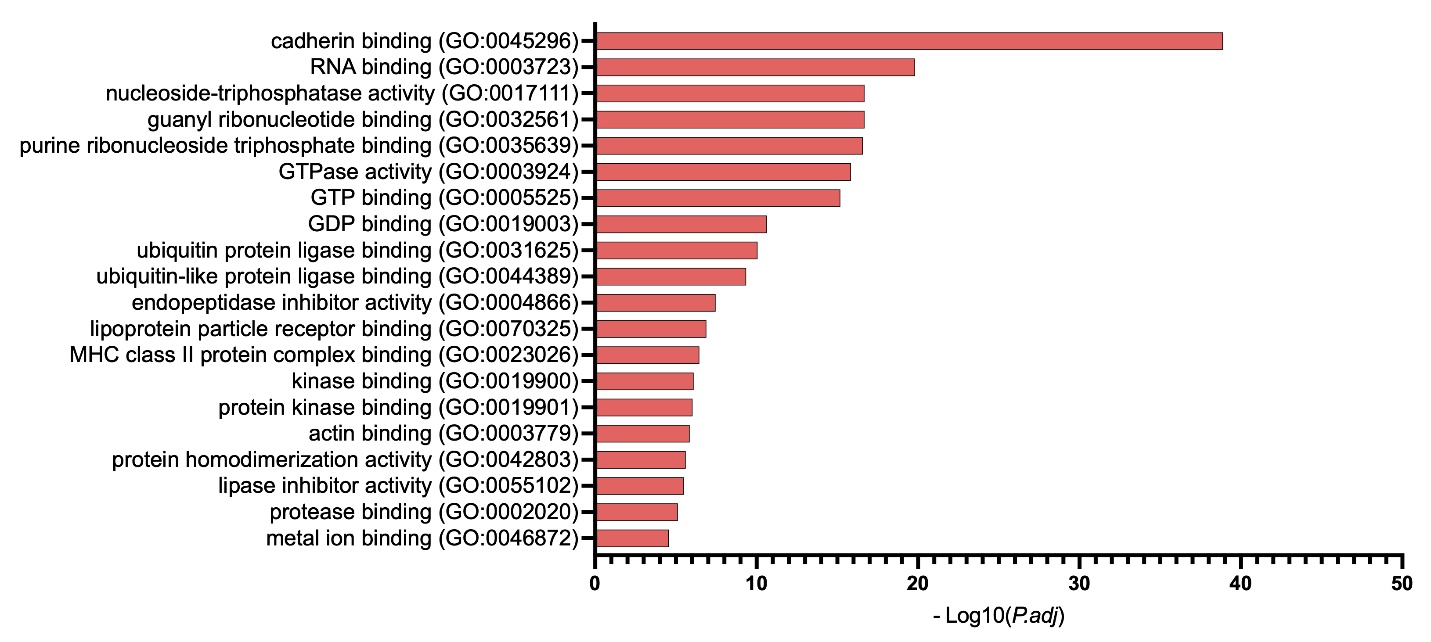

**Supplemental Figure 3. Top 20 Enriched Pathways Associated with Proteins in Naïve CEC-sEVs and m214-sEVs.**

Pathway enrichment analysis was performed using the Gene Ontology (GO) Molecular Function 2025 database to identify functional categories associated with proteins detected in CEC-sEVs and m214-sEVs. The top 20 pathways were ranked by statistical significance, expressed as −log₁₀ of the adjusted P-values. The raw data and results are listed in **Supplemental Excel 2.**

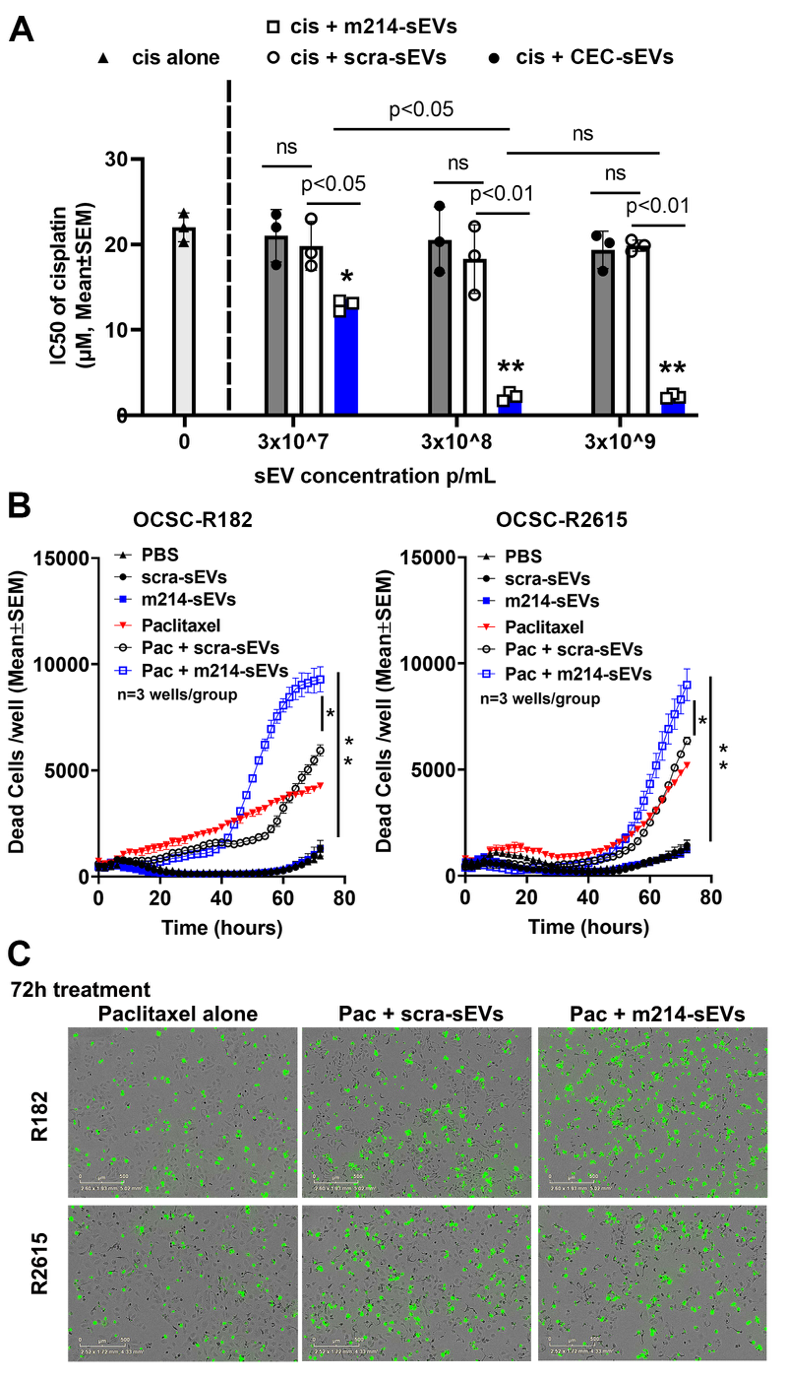

**Supplemental Figure 4. m214-sEVs Sensitize Chemoresistant OC Cells to Cisplatin and Paclitaxel.** (**A**) IC₅₀ values of cisplatin in A2780cis cells treated with increasing concentrations of sEVs (3×10⁷, 3×10⁸, or 3×10⁹ particles/mL) in combination with cisplatin. (**B**) Quantitative analysis of cytotoxicity in chemoresistant OCSC lines R182 and R2615, treated with paclitaxel (Pac; 20 µM) in combination with sEVs (3×10⁸ particles/mL) over 72 hours. Cell death was assessed using real-time CellTox Green assays. Data are presented as mean ± SEM from n = 3 wells per group. *p < 0.05, **p < 0.01 by one-way ANOVA with Tukey’s post hoc test. (**C**) Representative fluorescence images of OCSC-R182 and OCSC-R2615 cultures captured 72 hours post-treatment. Dead cells were indicated by green fluorescence. Treatment groups included paclitaxel alone, paclitaxel combined with scra-sEVs, and paclitaxel combined with m214-sEVs. Scale bars = 500 µm.

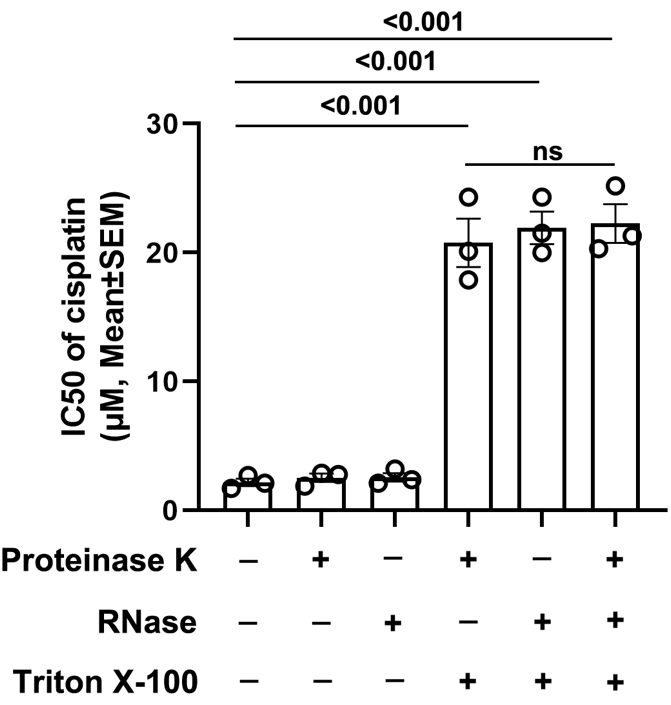

**Supplemental Figure 5. Vesicle Integrity Is Required for m214-sEV–Mediated Sensitization to Cisplatin**

The role of vesicle integrity in m214-sEV–mediated cisplatin sensitization was evaluated in A2780cis cells. m214-sEVs were pretreated with Proteinase K or RNase, with or without the membrane-disrupting detergent Triton X-100, prior to application on A2780cis cells. Subsequent measurement of cisplatin IC₅₀ values revealed that intact m214-sEVs significantly reduced cisplatin resistance.  This sensitizing effect was abolished when vesicle membranes were disrupted by Triton X-100, indicating that structural integrity is critical for functional delivery. Data represent mean ± SEM (n = 3).

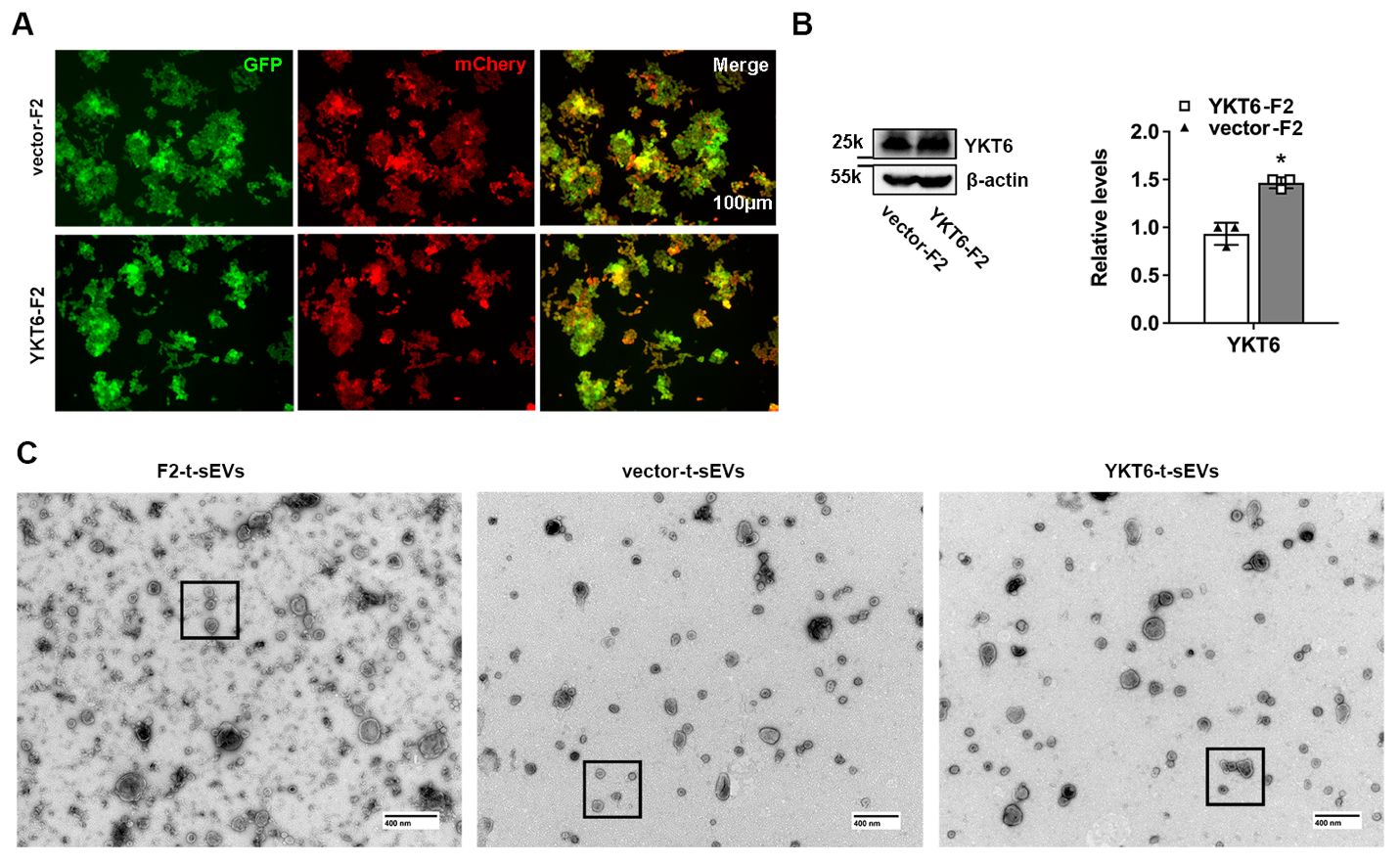

**Supplemental Figure 6. YKT6 Overexpression in OCSC1-F2 Cells and TEM Overview of Derived t-sEVs.**

(**A**) Representative fluorescence microscopy images showed co-expression of GFP (vector marker) and mCherry (cell label) in both vector-F2 and YKT6-F2 cells, confirming successful transduction. (**B**) Western blot analysis and corresponding quantification demonstrated significantly elevated YKT6 protein levels in YKT6-F2 cells compared to vector-F2 controls. *p < 0.05 by unpaired t-test. (**C**) Low-magnification TEM images showed the morphology of tumor-derived sEVs (t-sEVs) isolated from parental OCSC1-F2 cells, vector-F2 cells, and YKT6-F2 cells. Boxed areas corresponded to regions magnified in **Fig. 7H.**
